# Phenomic Versus Genomic Prediction - A Comparison of Prediction Accuracies for Grain Yield in Hard Winter Wheat Lines

**DOI:** 10.1101/2023.07.19.549748

**Authors:** Zachary J. Winn, Amanda L. Amsberry, Scott D. Haley, Noah D. DeWitt, R. Esten Mason

## Abstract

Common bread wheat (*Triticum aestivum* L.) is a key component of global diets, but the genetic improvement of wheat is not keeping pace with the growing demands of the world’s population. To increase efficiency and reduce costs, breeding programs are rapidly adopting the use of unoccupied aerial vehicles (UAVs) to conduct high-throughput spectral analyses. This study examined the effectiveness of multispectral indices in predicting grain yield compared to genomic prediction. Multispectral data were collected on advanced generation yield nursery trials during the 2019-2021 growing seasons in the Colorado State University Wheat Breeding Program. Genome-wide genotyping was performed on these advanced generations and all plots were harvested to measure grain yield. Two methods were used to predict grain yield: genomic estimated breeding values (GEBVs) generated by a genomic best linear unbiased prediction (gBLUP) model and phenomic phenotypic estimates (PPEs) using only spectral indices via multiple linear regression (MLR), k-nearest neighbors (KNN), and random forest (RF) models. In cross-validation, PPEs produced by MLR, KNN, and RF models had higher prediction accuracy (*r̄*: 0.41 ≤ *r̄* ≤ 0.48) than GEBVs produced by gBLUP (*r̄* = 0.35). In leave-one-year-out forward validation using only multispectral data for 2020 and 2021, PPEs from MLR and KNN models had higher prediction accuracy of grain yield than GEBVs of those same lines. These findings suggest that a limited number of spectra may produce PPEs that are more accurate than or equivalently accurate as GEBVs derived from gBLUP, and this method should be evaluated in earlier development material where sequencing is not feasible.

## INTRODUCTION

Common bread wheat (*Triticum aestivum* L.) is a diet staple across cultures and plays a key role in international food security. The global population is expected to rise to nearly ten billion people by the year 2050, yet annual genetic gain in wheat yield is not on track to meet this demand, which implies that a biocapacity deficit is inevitable unless significant action is taken (Alexandratos & Bruinsma, 2012; Berners-Lee et al., 2018; Ray et al., 2013). Genomic prediction is the process of using historical genomic and phenotypic data to produce genomic estimated breeding values (GEBVs) for lines which have only been sequenced and have not been evaluated via phenotyping (Meuwissen et al., 2001). Since the advent of high throughput genomic technologies, breeders have been using these predictions to implement genomic selection (GS) to increase selection intensity and heritability of selection in earlier generation material.

While genomic selection greatly assists breeders in making selections, it is not applied across all generations due to cost and labor limitations. High throughput spectral analysis via unoccupied aerial vehicles (UAVs) is becoming more common in modern breeding programs, generating large datasets of spectral reflectance and other phenotypic data. While the cost of collecting genome-wide markers on lines becomes prohibitive as the number of lines increases exponentially in earlier-generation trials, spectral data may be taken on all planted lines in all fields in a program at a fraction of the cost.

Recent research has suggested the utility of “phenomic selection” in breeding. Phenomic selection (PS) is the process of using electromagnetic spectral reflectance of organisms, captured by multispectral or hyperspectral sensors, to produce phenomic phenotypic estimates (PPEs) for traits of interest (Krause et al., 2019; Lane et al., 2020; Rincent et al., 2018; Robert et al., 2022; Sandhu et al., 2022). Recent publications have illustrated that spectral reflectance can create relationship matrices among individuals; furthermore, it has been shown that spectral reflectance matrices perform at or near the predictive ability of additive relationship matrices derived from either molecular markers or pedigree information when predicting grain yield (Krause et al., 2019). This may imply that phenomic selection could stand in place of genomic selection for selection candidates in earlier generations than those lines in the sequencing pipeline.

The potential application of PS in programs will be directly related to access of relevant technologies. Many of the publications on PS either rely on hyperspectral imaging derived from UAVs or near infrared (NIR) spectrometry taken in the field or in a lab setting (Krause et al., 2019; Lane et al., 2020; Rincent et al., 2018; Sandhu et al., 2022). These methods provide many different bands of reflectance to use for the creation of relationship matrices. However, their use within breeding programs may be limited due to the cost, labor, time, and the processing pipelines required of these techniques. Investigating if phenomic predictions may be made using a limited number of spectra as predictors on more accessible equipment may address this gap in application.

In the 2019, 2020, and 2021 growing seasons, the Colorado State University Wheat Breeding Program flew UAVs in cooperation with the Colorado State University Drone Center [https://www.research.colostate.edu/csudronecenter/] over fields containing late generation lines (F_3:5_, F_3:6_, and F_3:7+_) at the Agricultural Research, Development and Education Center (ARDEC; 4616 NE Frontage Road Fort Collins, CO 80524). Concurrently, those late development lines were sequenced for genome-wide markers. Using this unbalanced data set which models a true breeding system, the objectives of this study were to: (1) investigate the use of limited spectral information across years to produce PPEs, (2) compare calculated PPEs and GEBVs to adjusted means of yield, and (3) test several scenarios to observe which models produced optimal results for calculations of PPEs.

## MATERIALS AND METHODS

### Germplasm

The Colorado State University (CSU) Wheat Breeding Program pipeline is a modified bulk-pedigree method (Figure 1). All lines which occur in the later nurseries of the CSU breeding program are representative of the contemporary hard red and white winter wheat germplasm adapted to the targeted breeding areas of CSU; these areas include eastern Colorado, western Nebraska, western Kansas, and northwestern Oklahoma.

**Figure 1.**
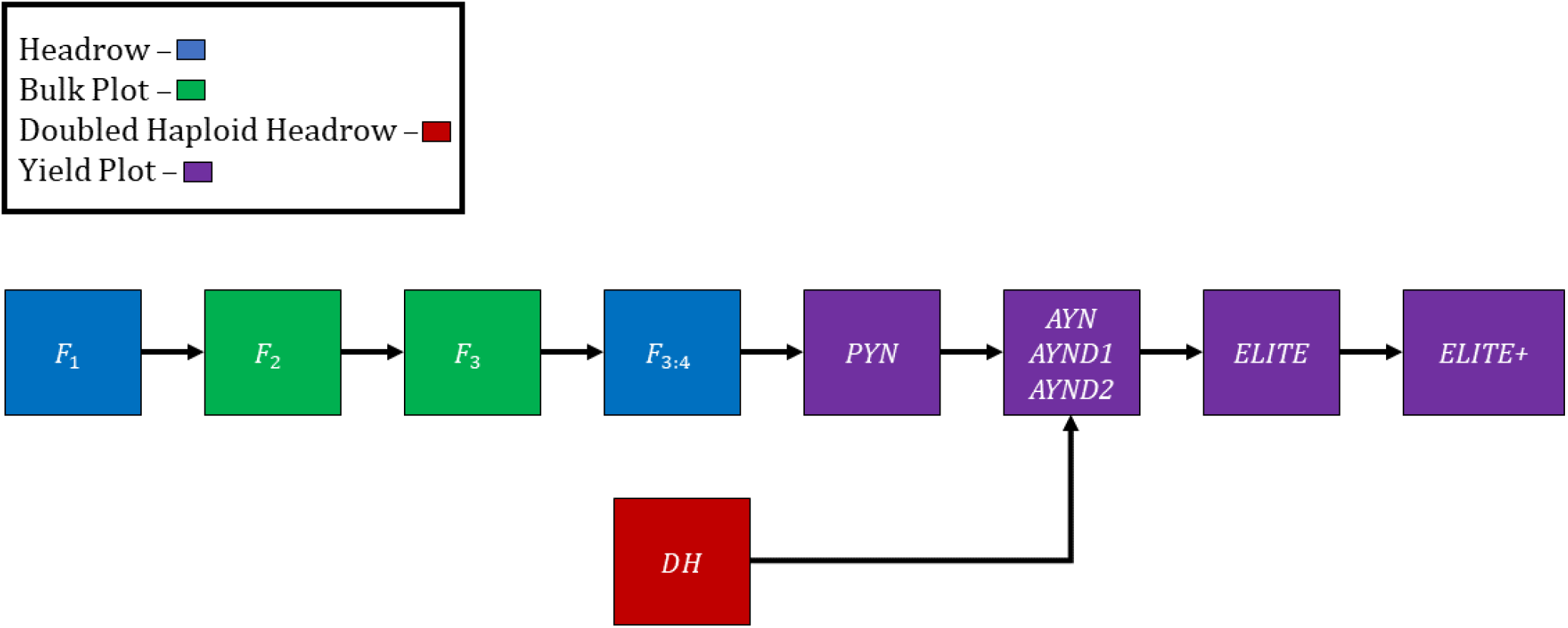
A diagram of the Colorado State University breeding program. Boxes represent germplasm/trials associated with the first few generations of inbreeding (F_1_ to F_3:4_), doubled haploids (DH), preliminary yield nursery (PYN), advanced yield nursery (AYN), double haploid advanced yield nursery one (AYND1) and two (AYND2), elite yield nursery (ELITE), and individuals that stay in the elite yield nursery for three to four years (ELITE+). Colors represent the plot type of each germplasm/trial. All generations after the F_3:4_ are F_3_-derived lines (*e.g.,* PYN = F_3:5_, AYN = F_3:6_, etc.).

The final single head selection occurs in an F_3_ bulk and selected heads are evaluated in rows in the following year. Final among-row selection occurs in the F_3:4_ and preliminary yield trials begin with F_3:5_ lines. The preliminary yield nursery (PYN; F_3:5_) is then followed by the advanced yield nursery (AYN; F_3:6_). All lines which make it past the PYN and AYN are then placed in the CSU Elite Trial (ELITE; F_5:7+_), where they remain until they are culled or released as a cultivar.

Since 2013, doubled haploids (DHs) have been routinely generated by the CSU Wheat Breeding Program via the F_1_ corn (*Zea maize* L.) pollen hybridization method (Santra et al., 2017). Once DH seed of a cross has been increased in a greenhouse, they are planted in rows and subjected to selection based on visual ratings and genomic predictions. Doubled haploid lines which are advanced from the row stage are then treated as AYN lines. These nurseries are conventionally titled AYND1 or AYND2 (“D” for doubled haploid) within a given year.

Spectral data were collected for three years of trials planted at the Agricultural Research, Development and Education Center (ARDEC) facility in Fort Collins, Colorado through collaboration with the CSU Drone Center (https://www.research.colostate.edu/csudronecenter/). Data from the following trials were used in the current work: the PYN from 2021; the AYN from 2019, 2020, and 2021; the AYND1 and AYND2 from 2019, 2020, and 2021, and the ELITE from 2019, 2020, and 2021. These trials resulted in a total dataset of 4,557 individual observations tied to a yield plot within a trial, within a year. A detailed description of the number of individuals in each nursery for the 2019, 2020, and 2021 season is provided (Table 1).

**Table 1:**
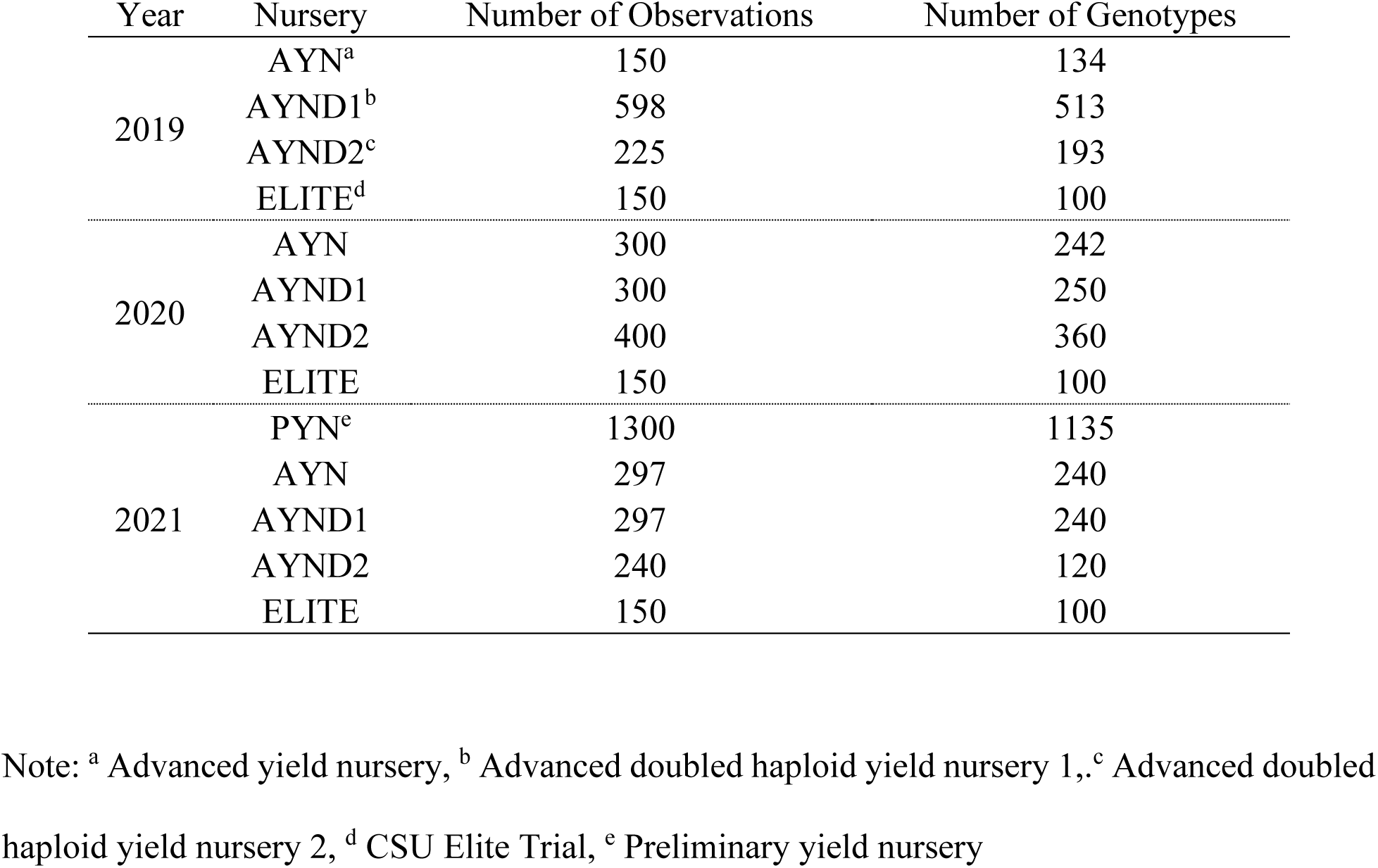
The number of individual genotypes per generation, per nursery, per year.

| Year | Nursery | Number of Observations | Number of Genotypes |
| --- | --- | --- | --- |
| 2019 | AYN<sup>a</sup> | 150 | 134 |
|  | AYND1<sup>b</sup> | 598 | 513 |
|  | AYND2<sup>c</sup> | 225 | 193 |
|  | ELITE<sup>d</sup> | 150 | 100 |
| 2020 | AYN | 300 | 242 |
|  | AYND1 | 300 | 250 |
|  | AYND2 | 400 | 360 |
|  | ELITE | 150 | 100 |
| 2021 | PYN<sup>e</sup> | 1300 | 1135 |
|  | AYN | 297 | 240 |
|  | AYND1 | 297 | 240 |
|  | AYND2 | 240 | 120 |
|  | ELITE | 150 | 100 |

Note: <sup>a</sup> Advanced yield nursery, <sup>b</sup> Advanced doubled haploid yield nursery 1, <sup>c</sup> Advanced doubled haploid yield nursery 2, <sup>d</sup> CSU Elite Trial, <sup>e</sup> Preliminary yield nursery

### Field Management and Phenotyping

All nurseries used in this study were planted at the ARDEC facility located in Fort Collins, Colorado at approximately 1,525 m above sea level. Within each growing season, all grain yield plots were planted in mid-September using a cable guided no-till drill seeder at 1.5m wide with 4.9m centers. During green up in the spring, plots were end trimmed using a 1.2m wide hooded sprayer and glyphosate (Monsanto, St Louis, Missouri, USA), leaving a harvestable area of 1.5m by 3.7m. All nurseries were planted in a statistical design which used row-column effects as blocking factors. The PYN was planted in an augmented block design; AYN, AYND1, and AYND2 trials were planted in an augmented block design; and ELITE trials were planted in a partially replicated design across all years.

Each year, soil samples were taken across fields to inform pre-plant fertilization regiments. In 2019, 64.61 kg ha^-1^ of 11-52-0 high phosphate MAP granular fertilizer with ammoniacal nitrogen was applied along with 349.83 kg ha^-1^ 46-0-0 urea granular nitrogen on September 6^th^, 2018, prior to planting. Post planting, 0.95 L ha^-1^ of Husky® (BASF; Ludwigshafen, Germany) was applied on April 24^th^, 2019 to control weeds. In 2020, 18.52 kg ha^-1^ of 0-0-62 high potassium potash was applied along with 32.63 kg ha^-1^ of 11-52-0 high phosphate MAP granular fertilizer with ammoniacal nitrogen and 358.99 kg ha^-1^ 46-0-0 urea granular nitrogen on September 13^th^, 2019, prior to planting. Post planting, 1.04 L ha^-1^ of Husky® was applied on April 28^th^, 2020 to control weeds. In 2021, After sampling to determine a proper fertilizing regiment, 32.62 kg ha^-1^ of 11-52-0 high phosphate MAP granular fertilizer with ammoniacal nitrogen was applied along with 358.18 kg ha^-1^ 46-0-0 urea granular nitrogen and 15.16 kg ha^-1^ of granular 90% sulfur on September 6^th^, 2020, prior to planting. Post planting, 1.04 L ha^-1^ of Husky® was applied on May 1^st^, 2021, to control weeds. Irrigation was applied, as needed as soon as temperatures allowed for irrigation lines to thaw in the spring. Prescriptive irrigation continued into grain fill and was terminated prior to physiological maturity and harvest. Detailed information on irrigation regiments is provided (Supplementary information 1).

Within each growing season, at physiological maturity, lines were harvested using a Zürn 150 (Zürn Harvesting GmbH and Company; Schöntal, Germany) research plot combine and measured for test weight, grain moisture, and grain yield using a HarvestMaster H2 Classic Graingage (Juniper Systems; Logan, Utah, USA). Grain yield was adjusted by moisture content so that the grain yields reflected 12% grain moisture, and yield was calculated using plot size metrics and reported in metric tons per hectare on a per-plot basis (t ha^-1^).

### Sequencing

For each line sequenced, ten seeds were planted, and a 2-3 cm tissue sample was taken from each plant and bulked together. Genomic quality deoxyribonucleic acid (DNA) was extracted using MagMax (ThermoFisher Scientific; Waltham, Massachusetts, USA) plant DNA kits. Extracted DNA was quantified using PicoGreen (ThermoFisher Scientific; Waltham, Massachusetts, USA) kits and normalized to a concentration of 20 ng µL^-1^ using an automated liquid handling system. Sequencing libraries were prepared according to Poland et al (2012).

All multiplexed libraries were sequenced on a NovaSeq 6000 (Illumina; San Diego, CA, USA) at 384-plex density per lane. Reads were aligned to the RefSeq Chinese Spring wheat reference sequence v2.0 using the burrow-wheeler aligner (Appels et al., 2018; Li & Durbin, 2009). Reads obtained were processed using the TASSEL 2.0 standalone pipeline and single nucleotide polymorphisms (SNPs) were organized into compressed variant calling format files (Danecek et al., 2011; Glaubitz et al., 2014). To produce full-rank genotypic matrices for genomic relationship matrix calculation, missing data were imputed using the Beagle algorithm V5.4 and the recombination distance-based map used for imputation was derived from a synthetic wheat biparental cross between ‘W7984’ and ‘Opata’ (Browning et al., 2018; Gutierrez-Gonzalez et al., 2019).

### Spectral Reflectance Acquisition

The processing pipeline for multispectral data consists of three distinct phases: spectral data collection via UAV-mounted cameras, image stitching and processing, and data extraction from individual plots. Between each year, flight parameters were changed at the discretion of the current drone technician to increase data quality, and this resulted in an unbalanced method of sampling from year-to-year. While individual processes and settings may have changed from year-to-year to improve the quality and efficiency of the pipeline over time, the general framework remained similar.

Drone flights were conducted at a height of approximately 120 m above ground level (AGL) in 2019, 100 m AGL in 2020, and 60 m AGL in 2021. This resulted in a resolution of 8.2 px cm^-1^ in 2019, 7.1 px cm^-1^ in 2020, and 4.2 px cm^-1^ in 2021 for the multispectral flights. The increased resolution of the images allowed for more data points per plot when calculating the averages for various spectral indices. In 2019 and 2020, ground control points (GCPs) in the form of plastic bucket lids were placed throughout the field including one at each corner, one at the midpoint of each edge, and at least one in the middle of the field. In 2021, no GCPs were used, which significantly increased the amount of manual post-flight processing.

Flights were conducted at the discretion of the current leader of the CSU Wheat Breeding Program and flights were taken when scheduling was available through the CSU Drone Center, leading to a wide variance among sampling dates and times. Over several flights taken between 2019-2022, the curated flight data has the narrowest window possible. A list of flight dates utilized in this study by year is provided (Table 2).

**Table 2.** Flight date within year.

| Year | Flight 1 | Flight 2 | Flight 3 | Flight 4 |
| --- | --- | --- | --- | --- |
| 2019 | 4/23/2019 | 5/24/2019 | 6/26/2019 | 7/12/2019 |
| 2020 | 5/1/2020 | 5/17/2020 | 6/7/2020 | 7/2/2020 |
| 2021 | 4/30/2021 | 5/15/2021 | 6/8/2021 | 6/18/2021 |
| Range | 8 days | 9 days | 19 days | 24 days |

Two different UAVs were used for flights. The primary UAV was a Matrice 600 Pro (DJI Technology; Nashan, Shenzhen, China) which housed two cameras – a MicaSense RedEdge-M (AgEagle Aerial Systems; Wichita, Kansas, USA) multispectral camera and a FLIR Duo Pro R (Teledyne FLIR; Wilsonville, Oregon, USA) thermal camera. The second UAV was a Phantom 4 RTK (DJI Technology; Nashan, Shenzhen, China) that gathered a set of backup RGB data. Between all sensors, data were collected for spectral bands representing blue [465-485 nm], green [550-570 nm], red [663-673 nm], red edge [712-722 nm], near-infrared (NIR) [820-860 nm], and long-wave infrared (LWIR or thermal) [7.5-13.5 μm].

All UAVs utilized real time kinematics (RTK) positioning for accurate geotagging of images. Images were collected in the nadir direction at +/-1 hour to solar noon. Immediately before and after each flight, the multispectral camera was calibrated using its accompanying calibration panel. The calibration panel used for each flight was a MicaSense RP04 model panel which had albedo values of 51.3 for blue at 475 nm, 51.3 for green at 560 nm, 51.3 for red at 668 nm, 51.2 for red edge at 717 nm, and 51 for NIR at 840 nm.

Pix4Dmapper (Pix4D SA; Prilly, Switzerland) was used to stitch the images together to create an orthomosaic for each of the spectral bands, including the visual bands collected with the RGB camera. These orthomosaics were then uploaded to a geodatabase in ArcGIS Pro (Esri; Redlands, California, USA) for further editing and orthorectifying. A single RGB image was selected as the image to georeference all other orthomosaics, which minimizes the geospatial error between orthomosaics and flights over the course of the growing season; this was done by georeferencing each orthomosaic’s GCPs to those visible in the reference RGB. In 2021, since no GCPs were placed in the field, other permanent, physical landmarks had to be used for reference.

Starting in 2021, excess green index (ExGI) was used as the source dataset for the ArcGIS machine learning classification function. The goal of using this feature was to remove the potential inaccuracy of the data in previous years by including soil surrounding the plot in the average spectral data calculations. The algorithm was run unsupervised with a high focus on grouping pixels based on both spatial proximity and spectral similarity. The algorithm was tasked with grouping the field into two self-identified categories using the ExGI orthomosaic for reference.

With only plants and soil present in the field, the algorithm successfully classified the field into these two groups. The result of this process was an additional raster layer in the geodatabase that specified boundaries around the plant matter in the field. Using these boundaries, the soil was cropped out of the orthomosaic images, ensuring that the only data remaining in the orthomosaics was related to plant matter. While retroactive processing of orthomosaics in 2019 and 2020 in this way is possible, resolution of previous flights was not conducive to separating soil from plant material, thus this procedure was only applied to 2021.

The average reflectance of each band was extracted per plot and ArcGIS Pro was used to calculate spectral indices including normalized difference vegetation index (NDVI). Sampling areas were defined using a Python script that retrieves field coordinates and dimensions from a CSV file and draws a polygon for each plot directly into the ArcGIS geodatabase that hosts the orthomosaics. Using the plot polygons as boundaries for the calculation of the averages, data tables containing the mean reflectance for each plot, date, band, and index were exported for use in further analysis.

### Validation Scenarios

Three separate scenarios were explored to assess how PPEs calculated for each line from plot-level spectral data compared to GEBVs and adjusted mean yield. In all scenarios presented, no genotypes were replicated between the training and test sets to avoid bias in the calculation of GEBVs or PPEs.

Scenario one (S1) involved taking the total available data across years, partitioning 80% of the available yield data to training across years and trials, and validating on the remaining 20%, while including all spectral data in both training and validation sets. This scenario most closely resembles a classic cross-validation scheme in genomic prediction. This process was replicated 30 times to obtain distributions of cross-validated accuracies.

Scenario two (S2) involved calculating adjusted means on a nursery-by-nursery basis and predicting yield in a trial where yield and spectral data from the same environment are in the training data. In S2 we assume that we have historical spectral and yield information on fields, and we have spectral and yield data on plot-level observations of other nurseries in the same field as a nursery which was not measured for yield. In S2, a single trial is left out of the data set and predicted using the rest of the available data. This may be applicable in a scenario where harvest of a trial is not feasible and grain yields must instead be predicted. This may be denoted as “hold-one-trial-out forward validation”.

Scenario three (S3) involved taking the total available data within two of the three available years, holding out yield data but not spectral data from that year, and using those data to train a model to predict the remaining year. For example, to predict the yield of all genotypes in 2021, both yield and spectral data from 2019 and 2020 and only spectral data from 2021 were used to train and predict yield data from 2021. This method implies that the program would have historical spectral data on a specific field, and that that historical data could be used to train a model to predict yield in an unobserved year for an unobserved genotype with spectral data but missing yield data. This scenario more closely resembles forward validation in genomic selection and may be denoted as “hold-one-year-out forward validation”.

### Statistical Analysis - Phenotypic Analysis and Genomic Prediction

All data management and statistical analysis was performed in R statistical software version 4.2.2 (R Core Team, 2022). All distributions for spectra and yield were checked for quality and normality using the qqplots function via the Base and Stats packages in R (R Core Team, 2022). All Q-Q plots exhibited near normality, warranting no transformation of any spectral indices or grain yield to fit normality. All mixed linear and genomic prediction models were performed using the package ASReml-R (Butler et al., 2009).

For calculation of adjusted means for both spectral indices and yield, the following univariate mixed linear model was used in every trial:

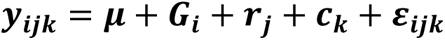

Where ***y*_*ijk*_** is the response, ***μ*** is the intercept, ***G*_*i*_** is the fixed genotypic response, ***r*_*j*_** is the random row effect, ***c*_*k*_** is the random column effect, and ***ε*_*ijk*_** is the residual where 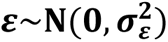. In each trial, random blocking effects for row and column could have either an identically and independently distributed (IID) covariance (*e.g.,* 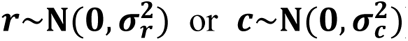) or covariance defined by autoregressive order one (AR1). In the case of autoregressive order one covariance, the covariance structure of either row or column random effects would appear as such:

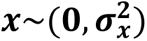

Where 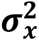 is equivalent to:

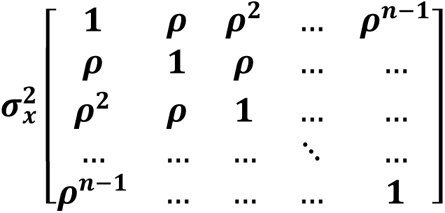

Where ***ρ*** represents the covariance between columns or rows and it is assumed that rows or columns further away from each other have a lower covariance.

Four total models (IID-column + IID-row, AR1-column + IID-row, IID-column + AR1-row, and AR1-column + AR1-row) were considered for each trial, and the optimal model was selected based on lowest achieved Bayesian information criterion (BIC). The resulting model with the lowest BIC was then used to produce adjusted means for the genotypic fixed effect for the subsequent model response.

To calculate adjusted means across trials within a year or across the total data set, the following mixed linear model was employed using the adjusted means calculated on a per-trial basis:

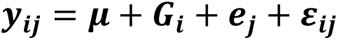

Where ***y*_*ij*_** is the response, ***μ*** is the intercept, ***G*_*i*_** is the fixed genotype effect, ***e*_*j*_** is the random environment effect 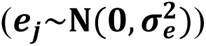, and ε_*ij*_ is the residual 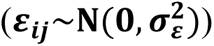. An “environment” in this case is the year that the trial was planted, concatenated to the name of the trial and the location of that trial (*e.g.,* “ARDEC_2019_AYN”). The adjusted means obtained across the entire dataset were used in S1, the adjusted means calculated with-in each trial were used in S2, and the adjusted means calculated within each year were used in S3.

For calculation of GEBVs in either S1, S2, or S3 the following genomic best linear unbiased prediction (gBLUP) model was employed:

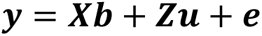

Where **y** is the response, ***X*** is the design matrix for fixed effects, ***b*** is the vector of fixed effects, ***Z*** is the design matrix for random genetic effects, and ***u*** is the vector of additive genetic effects defined by the genomic relationship matrix ***G*** 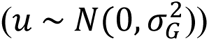 (VanRaden, 2008). Relationship matrix ***G*** was calculated from markers and weighted by allele frequency.

Genomic, narrow-sense, per-plot, heritabilities (*h^2^_g_*) were estimated for each trial in each year using the error and genotypic variances derived from gBLUP model. The gBLUP model used to estimate heritability is the same as presented above, however no random effects were fit for either row or column effect. Derived variances were then used to calculate 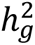 as such:

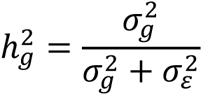

Where 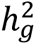 is genomic, narrow sense, per plot heritability; 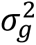 is genetic variance; and 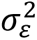 is the residual variance. This estimate of genomic, narrow sense, per plot heritability is derived from the genetic variance of a gBLUP model divided by the total variance. The genotypic variance was defined by the additive relationship matrix among individuals, and this estimate of heritability is an approximation of narrow-sense heritability, similar to estimates derived by population design, when all markers in linkage with causal effects are present in the genetic dataset (de Los Campos et al., 2015). Heritabilities were calculated using the vpredict function in ASReml-R (Butler et al., 2009).

### Statistical Analysis - Phenomic Prediction

Phenomic phenotypic estimates were calculated via multiple linear regression (MLR), k-nearest neighbors (KNN), and random forest (RF). All models used in calculation of PPEs were implemented using the package caret in R (Kuhn, 2008, 2022). In MLR, spectral indices were used as direct regressors on the response (adjusted means for grain yield). The MLR model may be visualized as:

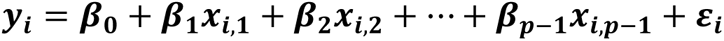

Where ***y*_*i*_** is the response, ***β*_0_** is the intercept multiplied by a constant of one, subsequent ***β_n_*** are the regression coefficients on spectral variables 1 through ***p***, ***p*** represents a spectral variable (a spectral index), and the residual error is IID 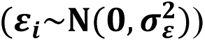. The estimate derived from the linear model on observations which only have spectral indices is interpreted as the PPE.

In the case of the nonparametric machine learning algorithms to follow, a seed was set for reproducibility in the case of S2 and S3, and 1000 permutations of five-fold cross validation in the training set was used to select the optimal hyper-parameters based on lowest achieved residual mean square error. When KNN models are used for regression of a continuous response, the data in a training set are used to construct a rule on how many nearest neighbors (*k*) to use in estimating the response in a test set. Estimation of the response is done directly by taking the *k* number of individuals in the training set most like the unobserved individual and directly averaging them to estimate the unobserved individual (Kramer & Kramer, 2013; Taunk et al., 2019).

In the KNN algorithm used in this study, tested values of *k* ranged from one to 50 in groups of five (*e.g., k* = 1, 5, 10,…,50). Based on cross-validation hyper-parameter tuning of the training data, we chose the value of *k* which minimized the residual mean square error. The average among the k-nearest neighbors used for estimating the unobserved individual in a validation set is interpreted as the PPE in this case.

Random forest algorithms are based on binary recursive partitioning to generate randomly designed decision trees. Random forest models can predict categorization by using these randomly generated decision trees to “vote” on categorical responses. However, in RF models used for regression, decision trees cannot take a direct voting approach when the response is continuous. Rather, RF models which are used for regression create cut-off points in the continuous response and use predictor variables (in this case, spectral reflectance data) to define an estimate of the response (grain yield) by averaging observations in the training data set that meet the same criterion.

For continuous predictor variables, finding the best possible split involves organizing the values of the predictors and identifying unique splits between every distinct pair of consecutive values. Normally, the midpoint of the interval between continuous response points is used as the point of division. These splits are then used to bifurcate the tree. In practice, many hundreds of trees are made using this method, and unobserved individuals are passed down these regression trees to form an estimation of the response. The prediction of the continuous response of the unobserved individual is equivalent to the summation of each estimation derived from each random regression tree in the random forest (Biau & Scornet, 2016; Breiman, 2001). This summation is interpreted as the PPE.

For the random forest model used in this study, defining variables (spectra) used for randomly splitting nodes in regression trees ranged from one to 20 in groups of 5 (*e.g.,* 1, 5, 10,…, 20). The number of random decision trees in the random forest was optimized using the caret R package. Based on cross validation hyper parameter tuning, we selected the *n* number of random splitting nodes that resulted in lowest residual mean square error in the training set.

To compare the predictive ability of GEBVs against PPEs, the GEBVs and PPEs were regressed on the adjusted means. Genomic and phenomic prediction accuracies were equivalent to the correlation of the corresponding GEBV or PPE to the adjusted means of grain yield. Both the Pearson’s correlation (*r*) and the coefficient of determination (*r^2^)* were reported as well as the residual mean square error (RMSE). In the current work, predictive ability is defined as the Pearson’s correlation coefficient between either the GEBVs or PPEs and the adjusted mean relevant to the scenario proposed.

## RESULTS

### Heritabilities and Correlations of Spectra and Grain Yield

In general, the genomic, narrow-sense, per-plot heritability of spectra was lower in 2019 and 2020 in comparison to the 2021 growing season (Figure 2). In the 2021 growing season, most of the ranges for estimated heritability of spectra did not include zero in their interval and had smaller ranges than in previous years. The mean, minimum, maximum, and standard deviation of all heritabilities observed within a year for grain yield and all spectra are provided (Supplemental Information 2).

**Figure 2.**
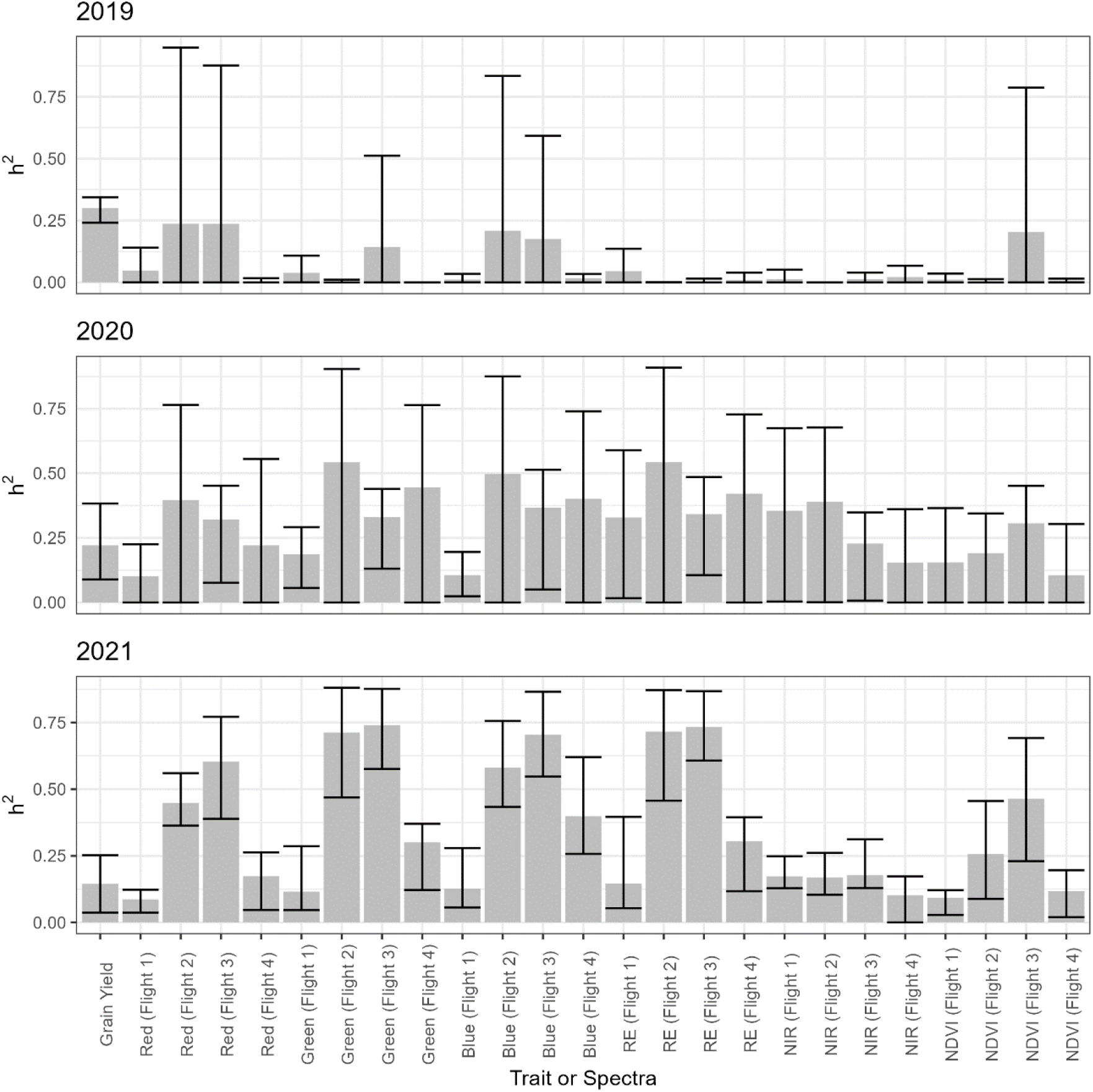
Bar-chart of average heritability across trials with each growing cycle. On the y-axis is the narrow-sense, per-plot, genomic heritability (h^2^). The x-axis depicts grain yield, red-blue-green (RGB) spectral reflectance, red edge (RE), near-infrared (NIR), and normalized difference vegetation index (NDVI). The flight from which each spectral index was collected is noted next to the spectrum name (*e.g.,* Red (Flight 1)). Each bar represents the average heritability of the corresponding trait on the x-axis. The black error lines surrounding each bar represent the observed range for that heritability across nurseries.

Correlation of spectral adjusted means to grain yield across trials within each year indicated that spectra were significantly correlated with grain yield in 2021 (Figure 3). However, correlations were, for the most part, sparsely correlated or near zero for the 2019 and 2020 growing seasons. In 2021, most spectral data (RGB, RE, and NIR) collected from later flights had significant negative correlations with grain yield, while NDVI alone had significant positive correlations to yield. The significant positive correlation of grain yield to NDVI implies that plots with higher photosynthetic activity tended to yield higher, a trend consistent with other research in wheat (Freeman et al., 2003; Hassan et al., 2019).

**Figure 3.**
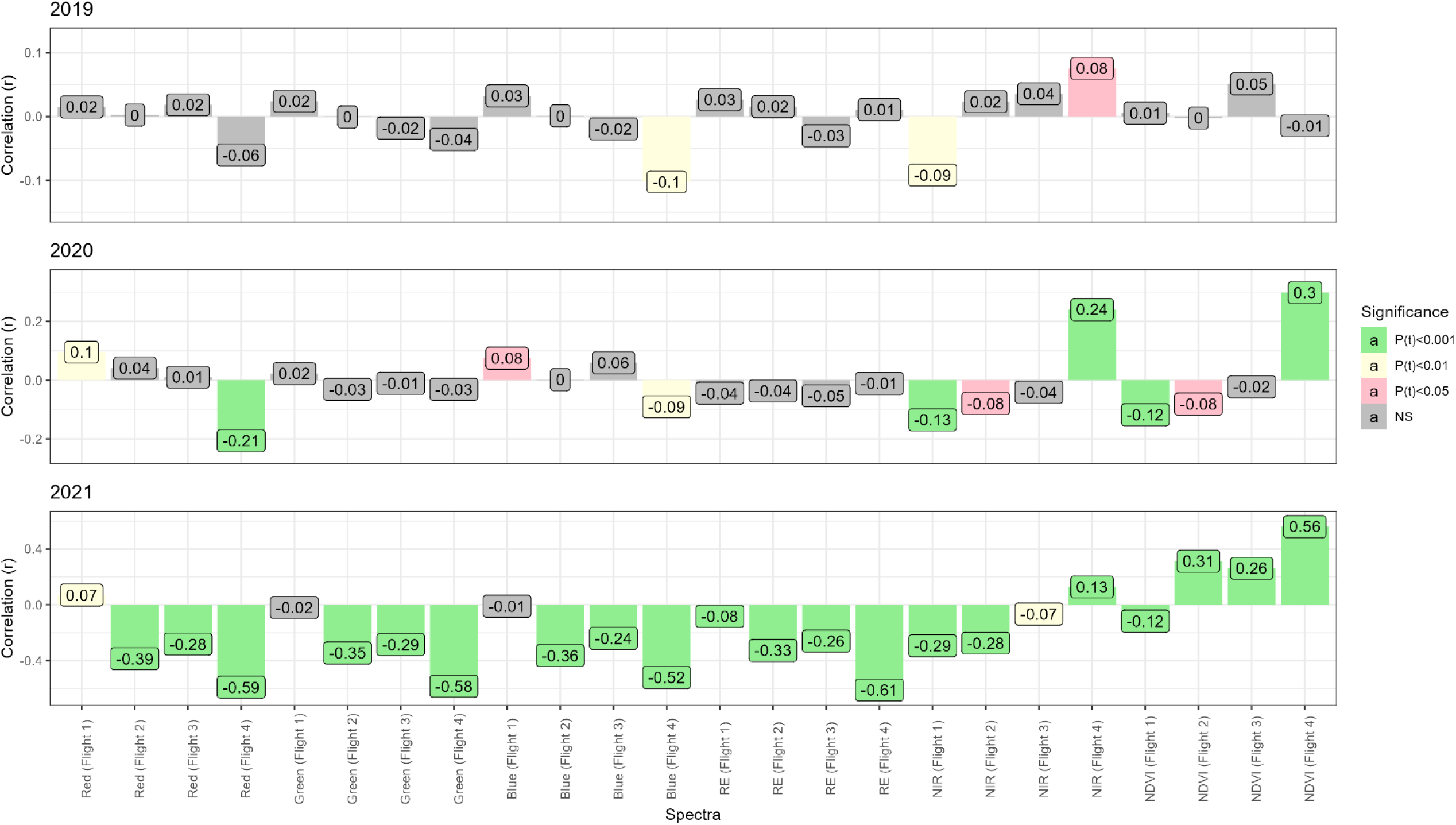
Bar-chart of Pearson’s correlation coefficient (r) of spectra vs grain yield. The x axis depicts different combinations of spectra and flights. Each bar represents the Pearson’s correlation coefficient of the corresponding trait on the x axis to grain yield. The color of each bar represents the significance of that correlation. The figure legend denoting significance can be found to the right of the graphs.

### Scenario 1: Cross-Validation

Randomly partitioned 80:20 training-test cross validation across 30 permutations revealed that gBLUP models, which only use additive genetic relationship information derived from molecular markers, underperformed in comparison to MLR, KNN, and RF models using only the five spectral indices recorded over four flights (Figure 4A). The average cross validated prediction accuracy of GEBVs derived from gBLUP was *r̄* = 0.35. By comparison, the average cross validated prediction accuracy of PPEs derived from the MLR, KNN, and RF models were *r̄* = 0.41, *r̄* = 0.44, and *r̄* = 0.48; respectively.

**Figure 4.**
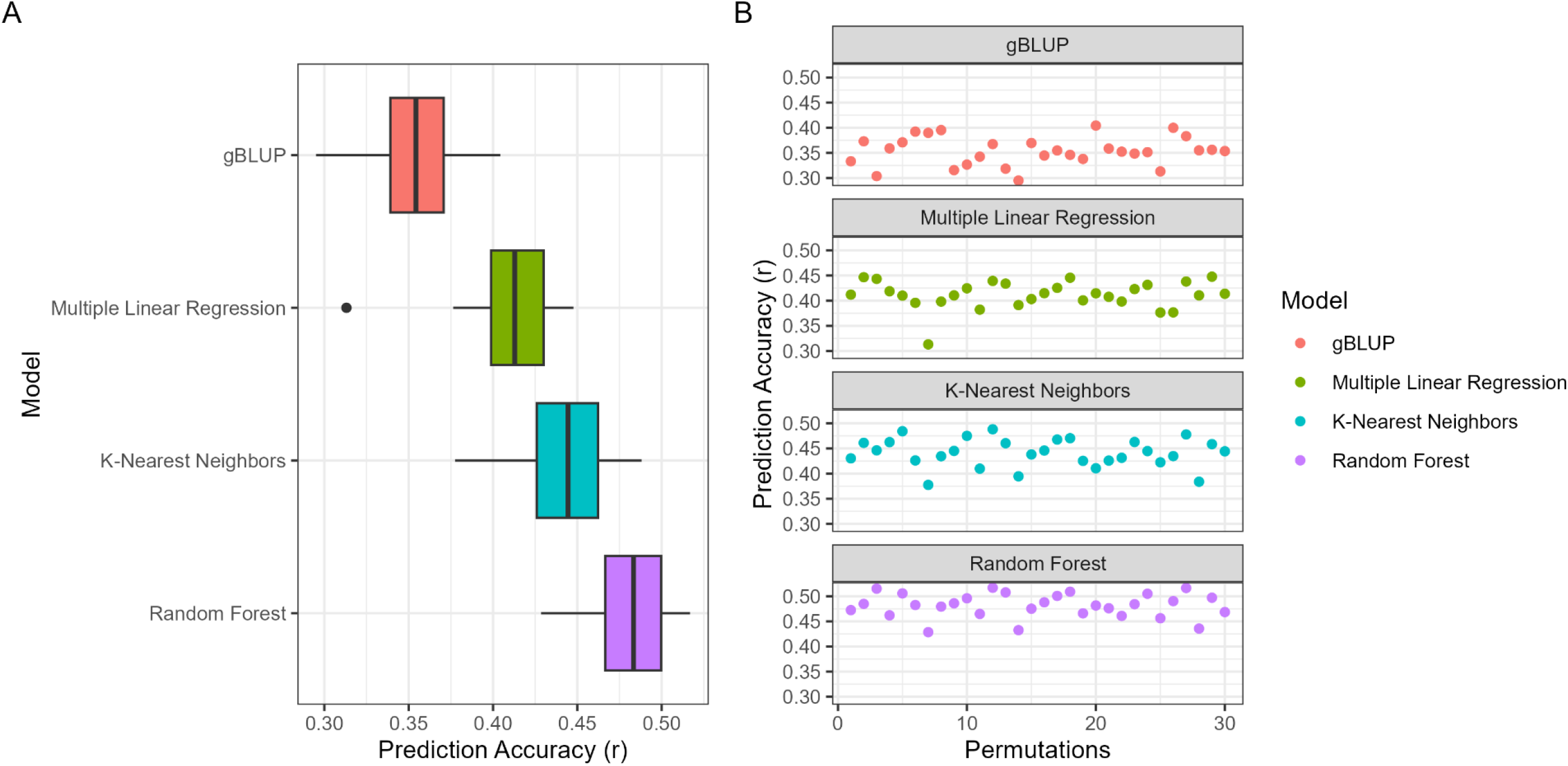
Boxplots of cross validation prediction accuracies **[A]** and a visualization of prediction accuracy (r) over thirty permutations **[B]**. On the y axis of panel **[A]** is the name of the model. On the x axis of panel **[A]** is the prediction accuracy. The sub-header of all graphs in panel **[B]** displays the name of the model. On the y axis of panel **[B]** is the prediction accuracy. On the x axis of panel **[B]** is the index for each permutation. The figure legend on the far right displays the corresponding colors for each model.

The standard deviation of the gBLUP, MLR, KNN, and RF predictions were 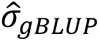 = 0.028, 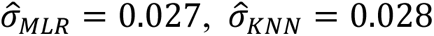, and 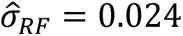; respectively. A visualization of the prediction accuracy per permutation is provided (Figure 4B). The RMSE for the gBLUP, MLR, KNN, and RF predictions were *RSME_gBLUP_* = 266.3, *RSME_MLR_* = 358.8, *RSME_KNN_* = 317.9, and *RSME_RF_* = 332.7; respectively. None of the models had noticeable differences in the standard deviation of their prediction accuracies, indicating that the range in prediction accuracies across permutations were similar. Of note, there was one prediction accuracy that significantly deviated in permutation seven of the MLR model (Figure 4B). A tabular summary of parameters from cross validation is also provided (Supplemental Information 3).

### Scenario 2: Leave-One-Trial-Out Forward Validation

In the 2019 season, PPEs underperformed in terms of prediction accuracy (*r*) in comparison to GEBVs (Figure 5). In the 2020 trials, the only cases where the prediction accuracies for PPEs were comparable to or higher than those derived from GEBVs were in the AYND1 and AYND2. The gBLUP prediction accuracy for the AYND1 in 2020 was *r* = 0.33, compared to *r* = 0.41 and *r* = 0.35, respectively. Likewise, in the AYND2 in 2020, the prediction accuracy for gBLUP was *r* = 0.26 while the accuracies for MLR and KNN were *r* = 0.35 and *r* = 0.25, respectively.

**Figure 5.**
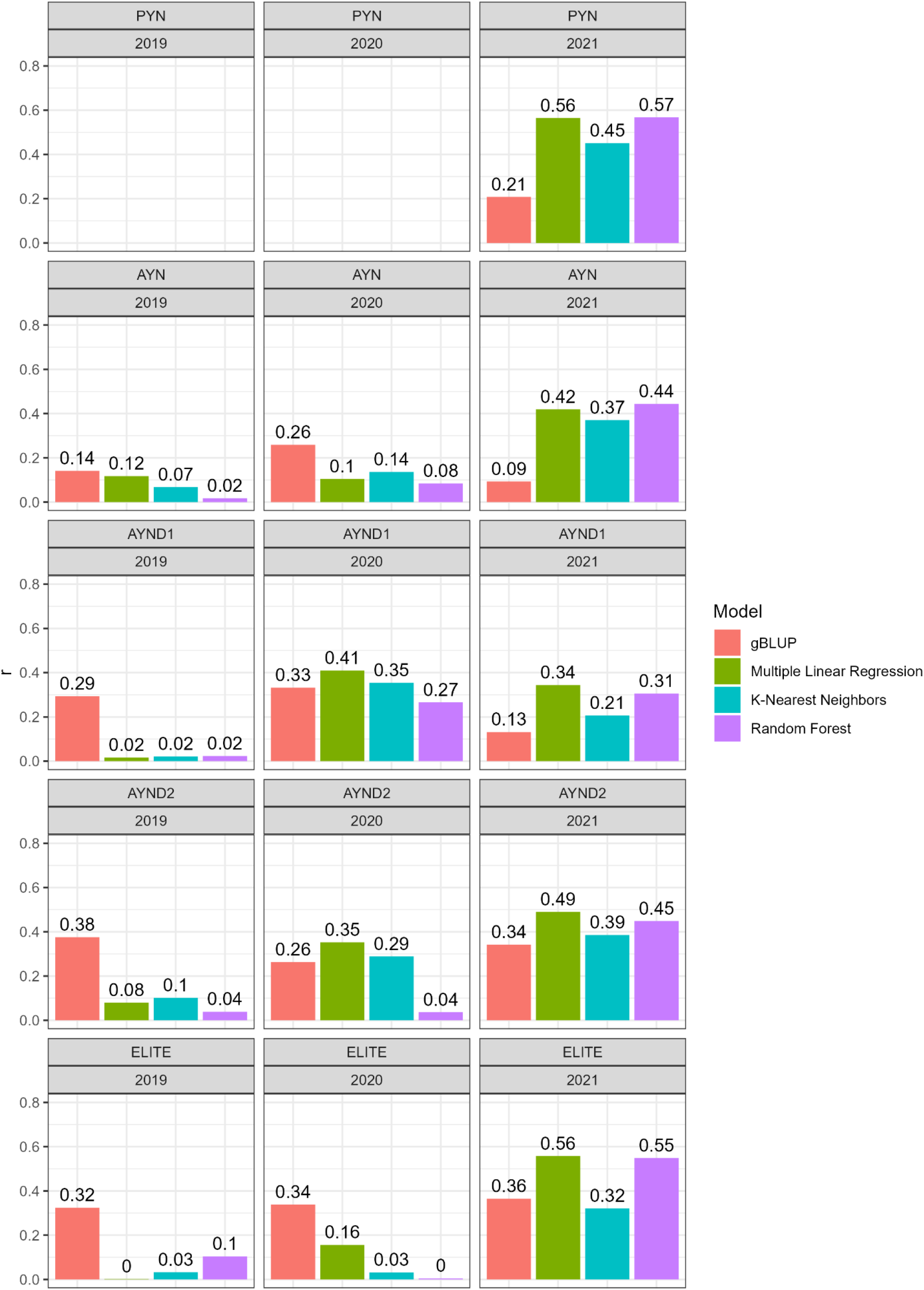
Bar graphs of leave-one-trial-out forward validation depicting the prediction accuracy (*r*) for each trial within each year. The y axis depicts the prediction accuracy as an expression of the Pearson’s correlation coefficient between the estimated breeding value and the best linear unbiased estimate for that trial. Each sub-graph is labeled by the trial to which the contents belong and the year that the trial was planted. Trial abbreviations are as follows: preliminary yield nursery (PYN), advanced yield nursery (AYN), advanced doubled haploid yield nursery one (AYND1), advanced doubled haploid yield nursery two (AYND2), and elite yield nursery (ELITE). Bars are colored to denote the model used. The number displayed above each bar represents the prediction accuracy of that model, in that year, for that trial. Subgraphs which do not contain content symbolize the unavailability of that data within that year (*e.g.,* PYN 2019 and 2020).

The trends observed in the 2019 and 2020 data were not apparent in the 2021 data. Overall, PPE prediction accuracies outperformed GEBV prediction accuracies. In the PYN of 2021, the predictive accuracy of the gBLUP model was *r* = 0.21; by comparison, MLR, KNN, and RF had substantially higher prediction accuracies (*r* = 0.56, *r* = 0.45, and *r* = 0.57; respectively). In the AYN, AYND1, AYND2, and ELITE; the prediction accuracies derived from PPE estimates either performed on-par with or better than the prediction accuracy of GEBV estimates (Figure 5). A table of all model statistics is provided as well (Supplemental Information 4).

### Scenario 3: Leave-One-Year-Out Forward Validation

Similar to S2, the prediction accuracies of PPEs in 2019 underperformed in comparison to GEBVs (*r_GEBV_* = 0.15, *r_PEBV_* ≤ 0.05; Figure 6). In both 2020 and 2021, PPEs derived from MLR and KNN models consistently outperformed GEBVs derived from gBLUP models. Only in 2021 did the RF model perform above the gBLUP model. In 2020, the following prediction accuracies were observed: *r* = 0.18 for gBLUP, *r* = 0.30 for MLR, *r* = 0.25 for KNN, and *r* = 0.01 for RF. In 2021, the following accuracies were observed: *r* = 0.12 for gBLUP, *r* = 0.32 for MLR, *r* = 0.22 for KNN, and *r* = 0.26 for RF.

**Figure 6.**
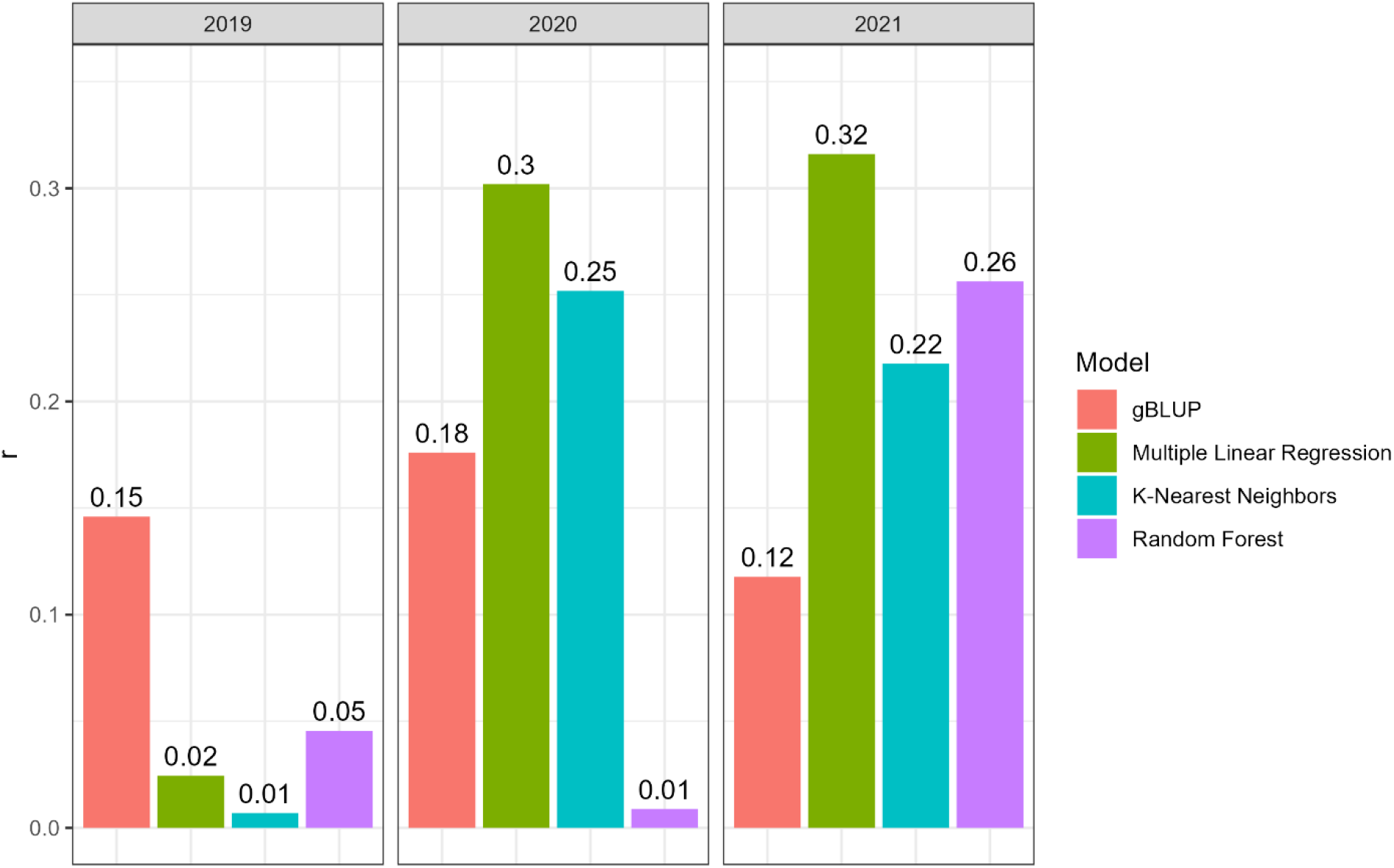
Bar graphs of leave-one-year-out forward validation prediction accuracies. The y axis depicts the prediction accuracy (*r*). Each subplot has a subtitle above it indicating the year to which the accuracy belongs. Each bar within each graph has been filled with a color that corresponds to what model that bar represents. The figure legend to the right denotes what model is represented by each color. Above each bar, in plain text, is the prediction accuracy.

## DISCUSSION

In the present work, genomic estimated breeding values were compared to predictions from spectral data (denoted as phenomic phenotypic estimates) through comparisons of prediction accuracies across several scenarios. In scenario one, we performed a cross validation study across thirty permutations and observed that models which produce PPEs using only five spectral indices over four flights were more accurate than GEBVs produced from gBLUP models. In scenario two, we saw that PPEs were more accurate than GEBVs when the heritability of the spectral indices was moderate to high and when spectra were significantly correlated with grain yield. In the third scenario, we saw that using data from different years to produce PPEs for an unobserved year produced more accurate results than GEBVs when heritability of spectral indices was moderate to high.

Rincent et al (2016) and Roberts et al (2022) suggest that spectral reflectance captures the endophenotypes of organisms and it is this phenomenon that allows for accurate predictions in PS. Furthermore, spectral reflectance can give estimates of plant health, disease pressure, and environmental stress (Du & Noguchi, 2017; Francesconi et al., 2021; Montesinos-López et al., 2017; Qin et al., 2022). This may suggest that PPEs include non-additive effects like environmental variation in their estimation, which may improve PPE model predictive accuracy.

Krause et al (2019) illustrated that relationship matrices can be calculated using hyperspectral reflectance and that those relationships derived from those matrices can be just as effective at predicting grain yield as relationship matrices derived from genetic data. Rincent et al (2016) found that wheat grain yield and heading date could be accurately predicted within and among environments using near-infrared spectroscopy to produce PPEs. Moreover, Krause et al (2019) reported that the level of prediction accuracy was highly correlated with the strength of relationship between hyperspectral reflectance and grain yield. Krause et al (2019) also found that heritability was heterogenous across timepoints and that this affected prediction accuracy, which is consistent with the current work’s results.

Montesinos-López et al (2017) found that removing ‘noisy bands’ of hyper-spectral reflectance which had heritability lower than *H* = 0.50 increased the predictive accuracy of their models for grain yield in wheat. Furthermore, Montesinos-López et al (2017) found that prediction accuracies were higher when they used all bands of spectra rather than one or a few vegetative indices. Rincent et al (2016) illustrated that accurate predictions for poplar tree (Genus *Populus*) bud flesh could also be achieved through phenomic prediction, meaning that this method could have practical application outside wheat. In support of that point, it was found by Lane et al (2020) that near-infrared spectroscopy readings from corn can accurately predict grain yield in environments under water stress and well-watered conditions.

In the present work, it was observed that the 2021 data showed higher heritability, higher correlations with grain yield, and improved prediction accuracies compared to those in 2019 and 2020. Two major differences in drone flights and data processing pipelines may be attributed to 2021 being the highest quality data in our dataset. Firstly, the reduction in flight height from 2019 to 2020, and then again in 2020 to 2021, resulted in higher image resolution with each subsequent year, which increased the amount of data points per plot in the calculation of average reflectance values. This most likely increased the accuracy of spectral averages derived from plots due to an increase in number of data points (pixels) per plot.

The second pipeline improvement which most-likely contributed to 2021 being the most informative data was the implementation of the excess green index to categorize and remove soil from the analysis in 2021. This process implemented in 2021 eliminated the risk of soil within the plot area skewing the averages away from representative plant reflectance values. While this process of classifying and removing soil could have been retroactively performed in previous years for a direct comparison, the flight heights in previous years led to a reduction in resolution, which hinders the programs’ ability to discriminate between soil and plant matter accurately. Therefore, we recommend reduction of flight heights to approximately 60m or less to increase resolution and the use of a similar machine learning method to remove non-plant material from images prior to extraction of spectra from images.

Another contributing factor to varying results across years may be related to flight dates (Table 2). In the current analysis, we took existing drone data taken within the CSU breeding program and aligned flight dates that were closest to each other in time. This data, while apparently informative, is not balanced. Moreover, growth stage in wheat is known to affect spectral reflectance (Sun et al., 2010; Xie et al., 2020). Therefore, to reduce variability in the dataset and potentially improve reproducibility of results, it is suggested to assess fields at the same growth stage across years.

In scenario one, where we compared 80:20 split cross-validation of GEBVs vs PPEs, we observed that all methods of estimation of PPEs were superior in their prediction accuracies in comparison to GEBVs. Thus, it appears that PPEs have similar predictive ability to GEBVs across years. However, it is important to note that the GEBVs estimated by our gBLUP models are based on additive effects, whereas the PPEs most-likely include environmental and non-additive effects in their calculation. Therefore, it would be important to conduct follow up studies on response to selection for PPEs vs GEBVs in early generation material.

While scenario one demonstrates the predictability of the dataset upon itself, scenario two, which involved predicting yield for plots where harvest is not possible, may have some direct utility in breeding programs. These trials in scenario two, which would be otherwise abandoned due to harvest conditions or labor constraints, may be flown with drones and predictions can be made without genetic information. This information could be used for making selections in limited datasets as well as supplementing training population data in genomic prediction by increasing the number of total observations for the phenotype of interest, yet these applications must be vetted in further studies.

Although scenario one and two yielded fascinating results, the scenario with the highest utility is scenario three, where we attempted to predict observations in an unobserved year with historical training data. While it appears that performing this procedure in years where spectral data collection is suboptimal (E.G., 2019) leads to underperformance in comparison to GEBVs (Figure 6), we did observe that in following years where spectral data was more heritable and better correlated to yield, that PPEs were comparable to or better than GEBVs. This may indicate that PPEs may be made on unobserved genotypes in unobserved years so long as the criteria of correlation and heritability are met.

A major component of the cost and labor of phenotyping trials for yield data involves the planting and harvesting of plots. Calculation of PPEs for this scenario three may allow breeders to plant larger trials of early-generation material and selectively harvest a subset of lines based on collection of spectral data. Although it is possible to use spectral data from plots prior to harvest to produce PPEs, this method only saves effort by allowing for selective harvest, and the same amount of financial and work resources would still need to be invested in planting. In contrast, genomic selection offers a way to reduce planting by using genotyping out of field and prediction, which can be more cost effective over time.

Selective harvest of plots can be challenging with mechanized harvest methods, and this may be less than optimal for programs which do not selectively harvest preliminary yield trials. In lieu of applying this method in the yield plot stage, perhaps the method proposed in scenario three would be best applied in the earlier generations where the number of individuals to genotype would be far too large for genomic selection and plots are harvest by hand.

Hypothetically, single plants or headrows of selection candidates could be planted with a subset of lines (a training population) which have replicated yield observations over years. The spectral information and historical yield data of the training population could then be used to produce PPEs for single plants or headrows which only have spectral data. This method is similar to the one proposed by Roberts et al (2022). Regardless, this method has yet to be tested for within-family prediction/selection and the response to phenomic selection in comparison to visual or genomic selection has not been reported in the literature.

The results of the current work, in correspondence with previously written literature, support the hypothesis that PPEs can be produced from spectra and that PPE accuracies are comparable to or better than GEBV accuracies, if the spectra are heritable and are correlated with the trait of interest. More interestingly, we have demonstrated that a limited number of spectral bands, rather than the wide range of spectra derived from hyperspectral analysis, may be used to create comparable predictions.

The presented results indicate that the 2019 spectral data was suboptimal due to sampling methods; however, models in scenario 3, which included the 2019 and 2020 spectral data to predict grain yield in 2021, managed to produce prediction accuracies higher than that of gBLUP (Figure 6), which seems to suggest that large datasets over time can compensate for suboptimal years of data collection. Thus, with further investigation and restructuring of breeding methods, phenomic prediction may be an effective alternative to genomic prediction in earlier generation material.

## ACKNOWLEDGMENTS

This research was made possible by funds derived from the competitive grant 2022-68013-36439 (WheatCAP) from the USDA National Institute of Food and Agriculture.

## SUPPLEMENTAL MATERIAL

**Supplemental Information 1.** Watering regiment by season.

| Growing Season | Date | Amount of Water (cm) |
| --- | --- | --- |
| 2019 | 6/26/2019 | 3.18 |
| 2019 | 6/13/2019 | 2.54 |
| 2020 | 6/25/2020 | 2.54 |
| 2020 | 6/18/2020 | 2.54 |
| 2020 | 6/12/2020 | 1.27 |
| 2020 | 6/4/2020 | 3.18 |
| 2020 | 5/28/2020 | 3.05 |
| 2020 | 5/22/2020 | 2.54 |
| 2020 | 5/13/2020 | 3.18 |
| 2020 | 4/7/2020 | 2.16 |
| 2021 | 6/3/2021 | 3.18 |
| 2021 | 6/13/2021 | 3.30 |
| 2021 | 6/18/2021 | 3.18 |
| 2021 | 6/24/2021 | 3.18 |

**Supplemental Information 2.** Average heritabilities within each growing season.

| Year | Trait | $h^2$ | | | |
| --- | --- | --- | --- | --- | --- |
|  |  | Mean | Min | Max | SD |
| 2019 | Grain Yield | 0.30 | 0.24 | 0.34 | 0.04 |
|  | Red (Flight 1) | 0.05 | <0.01 | 0.14 | 0.06 |
|  | Red (Flight 2) | 0.24 | <0.01 | 0.95 | 0.47 |
|  | Red (Flight 3) | 0.24 | <0.01 | 0.88 | 0.43 |
|  | Red (Flight 4) | <0.01 | <0.01 | 0.02 | <0.01 |
|  | Green (Flight 1) | 0.04 | <0.01 | 0.11 | 0.05 |
|  | Green (Flight 2) | <0.01 | <0.01 | 0.01 | <0.01 |
|  | Green (Flight 3) | 0.14 | <0.01 | 0.51 | 0.25 |
|  | Green (Flight 4) | <0.01 | <0.01 | <0.01 | <0.01 |
|  | Blue (Flight 1) | 0.01 | <0.01 | 0.03 | 0.02 |
|  | Blue (Flight 2) | 0.21 | <0.01 | 0.83 | 0.42 |
|  | Blue (Flight 3) | 0.18 | <0.01 | 0.59 | 0.28 |
|  | Blue (Flight 4) | 0.02 | <0.01 | 0.03 | 0.01 |
|  | RE (Flight 1) | 0.05 | <0.01 | 0.14 | 0.06 |
|  | RE (Flight 2) | <0.01 | <0.01 | <0.01 | <0.01 |
|  | RE (Flight 3) | <0.01 | <0.01 | 0.02 | <0.01 |
|  | RE (Flight 4) | 0.01 | <0.01 | 0.04 | 0.02 |
|  | NIR (Flight 1) | 0.01 | <0.01 | 0.05 | 0.03 |
|  | NIR (Flight 2) | <0.01 | <0.01 | <0.01 | <0.01 |
|  | NIR (Flight 3) | 0.01 | <0.01 | 0.04 | 0.02 |
|  | NIR (Flight 4) | 0.02 | <0.01 | 0.07 | 0.03 |
|  | NDVI (Flight 1) | 0.01 | <0.01 | 0.04 | 0.02 |
|  | NDVI (Flight 2) | <0.01 | <0.01 | 0.01 | <0.01 |
|  | NDVI (Flight 3) | 0.20 | <0.01 | 0.79 | 0.39 |
|  | NDVI (Flight 4) | <0.01 | <0.01 | 0.02 | <0.01 |
| 2020 | Grain Yield | 0.22 | 0.09 | 0.38 | 0.12 |
|  | Red (Flight 1) | 0.10 | <0.01 | 0.23 | 0.09 |
|  | Red (Flight 2) | 0.40 | <0.01 | 0.76 | 0.34 |
|  | Red (Flight 3) | 0.32 | 0.08 | 0.45 | 0.17 |
|  | Red (Flight 4) | 0.22 | <0.01 | 0.56 | 0.24 |
|  | Green (Flight 1) | 0.19 | 0.06 | 0.29 | 0.11 |
|  | Green (Flight 2) | 0.54 | <0.01 | 0.90 | 0.41 |
|  | Green (Flight 3) | 0.33 | 0.13 | 0.44 | 0.14 |
|  | Green (Flight 4) | 0.45 | <0.01 | 0.76 | 0.37 |
|  | Blue (Flight 1) | 0.11 | 0.02 | 0.20 | 0.07 |
|  | Blue (Flight 2) | 0.50 | <0.01 | 0.87 | 0.39 |
|  | Blue (Flight 3) | 0.37 | 0.05 | 0.51 | 0.22 |
|  | Blue (Flight 4) | 0.40 | <0.01 | 0.74 | 0.33 |
|  | RE (Flight 1) | 0.33 | 0.02 | 0.59 | 0.27 |
|  | RE (Flight 2) | 0.54 | <0.01 | 0.91 | 0.41 |
|  | RE (Flight 3) | 0.34 | 0.11 | 0.49 | 0.16 |
|  | RE (Flight 4) | 0.42 | <0.01 | 0.73 | 0.34 |
|  | NIR (Flight 1) | 0.35 | <0.01 | 0.67 | 0.33 |
|  | NIR (Flight 2) | 0.39 | <0.01 | 0.68 | 0.33 |
|  | NIR (Flight 3) | 0.23 | <0.01 | 0.35 | 0.15 |
|  | NIR (Flight 4) | 0.15 | <0.01 | 0.36 | 0.15 |
|  | NDVI (Flight 1) | 0.15 | <0.01 | 0.37 | 0.16 |
|  | NDVI (Flight 2) | 0.19 | <0.01 | 0.34 | 0.14 |
|  | NDVI (Flight 3) | 0.31 | <0.01 | 0.45 | 0.21 |
|  | NDVI (Flight 4) | 0.11 | <0.01 | 0.30 | 0.14 |
| 2021 | Grain Yield | 0.15 | 0.04 | 0.25 | 0.10 |
|  | Red (Flight 1) | 0.09 | 0.04 | 0.12 | 0.03 |
|  | Red (Flight 2) | 0.45 | 0.36 | 0.56 | 0.08 |
|  | Red (Flight 3) | 0.60 | 0.39 | 0.77 | 0.17 |
|  | Red (Flight 4) | 0.17 | 0.05 | 0.26 | 0.09 |
|  | Green (Flight 1) | 0.12 | 0.05 | 0.29 | 0.10 |
|  | Green (Flight 2) | 0.71 | 0.47 | 0.88 | 0.15 |
|  | Green (Flight 3) | 0.74 | 0.58 | 0.88 | 0.13 |
|  | Green (Flight 4) | 0.30 | 0.12 | 0.37 | 0.10 |
|  | Blue (Flight 1) | 0.13 | 0.06 | 0.28 | 0.09 |
|  | Blue (Flight 2) | 0.58 | 0.43 | 0.76 | 0.12 |
|  | Blue (Flight 3) | 0.70 | 0.55 | 0.87 | 0.13 |
|  | Blue (Flight 4) | 0.40 | 0.26 | 0.62 | 0.14 |
|  | RE (Flight 1) | 0.15 | 0.05 | 0.40 | 0.14 |
|  | RE (Flight 2) | 0.72 | 0.46 | 0.87 | 0.15 |
|  | RE (Flight 3) | 0.73 | 0.61 | 0.87 | 0.12 |
|  | RE (Flight 4) | 0.30 | 0.12 | 0.39 | 0.11 |
|  | NIR (Flight 1) | 0.17 | 0.13 | 0.25 | 0.05 |
|  | NIR (Flight 2) | 0.17 | 0.10 | 0.26 | 0.06 |
|  | NIR (Flight 3) | 0.18 | 0.13 | 0.31 | 0.08 |
|  | NIR (Flight 4) | 0.10 | <0.01 | 0.17 | 0.07 |
|  | NDVI (Flight 1) | 0.09 | 0.03 | 0.12 | 0.04 |
|  | NDVI (Flight 2) | 0.26 | 0.09 | 0.46 | 0.14 |
|  | NDVI (Flight 3) | 0.46 | 0.23 | 0.69 | 0.20 |
|  | NDVI (Flight 4) | 0.12 | 0.02 | 0.20 | 0.06 |

**Supplemental Information 3.** Summary of cross validation model performance.

| Model | r |  |  |  | r<sup>2</sup> |  |  |  | Residual Mean Square Error |  |  |  |
| --- | --- | --- | --- | --- | --- | --- | --- | --- | --- | --- | --- | --- |
|  | Min | Max | Mean | SD | Min | Max | Mean | SD | Min | Max | Mean | SD |
| gBLUP | 0.30 | 0.40 | 0.35 | 0.03 | 0.09 | 0.16 | 0.13 | 0.02 | 239.91 | 290.95 | 266.35 | 12.57 |
| Multiple Linear Regression | 0.31 | 0.45 | 0.41 | 0.03 | 0.10 | 0.20 | 0.17 | 0.02 | 297.01 | 358.86 | 323.03 | 13.50 |
| K-Nearest Neighbors | 0.38 | 0.49 | 0.44 | 0.03 | 0.14 | 0.24 | 0.20 | 0.02 | 285.92 | 345.02 | 317.94 | 15.10 |
| Random Forest | 0.43 | 0.52 | 0.48 | 0.02 | 0.18 | 0.27 | 0.23 | 0.02 | 307.13 | 349.75 | 332.70 | 11.96 |

**Supplemental Information 4.** Summary of year, trial, and model prediction accuracy (r), coefficient of determination (r^2^), and the residual mean square error (RMSE).

| Year | Trial | Model | r | $r^2$ | RMSE |
| --- | --- | --- | --- | --- | --- |
| 2019 | AYN | gBLUP | 0.14 | 0.02 | 238.94 |
| 2019 | AYN | Multiple Linear Regression | 0.12 | 0.01 | 397.19 |
| 2019 | AYN | K-Nearest Neighbors | 0.07 | 0.00 | 309.30 |
| 2019 | AYN | Random Forest | 0.02 | 0.00 | 248.40 |
| 2019 | AYND1 | gBLUP | 0.29 | 0.09 | 272.50 |
| 2019 | AYND1 | Multiple Linear Regression | 0.02 | 0.00 | 498.23 |
| 2019 | AYND1 | K-Nearest Neighbors | 0.02 | 0.00 | 216.75 |
| 2019 | AYND1 | Random Forest | 0.02 | 0.00 | 166.71 |
| 2019 | AYND2 | gBLUP | 0.38 | 0.14 | 331.15 |
| 2019 | AYND2 | Multiple Linear Regression | 0.08 | 0.01 | 396.67 |
| 2019 | AYND2 | K-Nearest Neighbors | 0.10 | 0.01 | 205.81 |
| 2019 | AYND2 | Random Forest | 0.04 | 0.00 | 187.52 |
| 2019 | ELITE | gBLUP | 0.32 | 0.11 | 238.04 |
| 2019 | ELITE | Multiple Linear Regression | 0.00 | 0.00 | 293.17 |
| 2019 | ELITE | K-Nearest Neighbors | 0.03 | 0.00 | 217.46 |
| 2019 | ELITE | Random Forest | 0.10 | 0.01 | 244.35 |
| 2020 | AYN | gBLUP | 0.26 | 0.07 | 259.32 |
| 2020 | AYN | Multiple Linear Regression | 0.10 | 0.01 | 291.18 |
| 2020 | AYN | K-Nearest Neighbors | 0.14 | 0.02 | 228.72 |
| 2020 | AYN | Random Forest | 0.08 | 0.01 | 267.27 |
| 2020 | AYND1 | gBLUP | 0.33 | 0.11 | 280.49 |
| 2020 | AYND1 | Multiple Linear Regression | 0.41 | 0.17 | 228.50 |
| 2020 | AYND1 | K-Nearest Neighbors | 0.35 | 0.13 | 132.37 |
| 2020 | AYND1 | Random Forest | 0.27 | 0.07 | 135.43 |
| 2020 | AYND2 | gBLUP | 0.26 | 0.07 | 250.53 |
| 2020 | AYND2 | Multiple Linear Regression | 0.35 | 0.12 | 362.54 |
| 2020 | AYND2 | K-Nearest Neighbors | 0.29 | 0.08 | 187.31 |
| 2020 | AYND2 | Random Forest | 0.04 | 0.00 | 169.65 |
| 2020 | ELITE | gBLUP | 0.34 | 0.11 | 243.18 |
| 2020 | ELITE | Multiple Linear Regression | 0.16 | 0.02 | 242.90 |
| 2020 | ELITE | K-Nearest Neighbors | 0.03 | 0.00 | 211.85 |
| 2020 | ELITE | Random Forest | 0.00 | 0.00 | 174.98 |
| 2021 | PYN | gBLUP | 0.21 | 0.04 | 164.62 |
| 2021 | PYN | Multiple Linear Regression | 0.56 | 0.32 | 245.68 |
| 2021 | PYN | K-Nearest Neighbors | 0.45 | 0.20 | 238.89 |
| 2021 | PYN | Random Forest | 0.57 | 0.32 | 280.31 |
| 2021 | AYN | gBLUP | 0.09 | 0.01 | 260.78 |
| 2021 | AYN | Multiple Linear Regression | 0.42 | 0.18 | 202.12 |
| 2021 | AYN | K-Nearest Neighbors | 0.37 | 0.14 | 155.51 |
| 2021 | AYN | Random Forest | 0.44 | 0.20 | 185.24 |
| 2021 | AYND1 | gBLUP | 0.13 | 0.02 | 260.14 |
| 2021 | AYND1 | Multiple Linear Regression | 0.34 | 0.12 | 249.78 |
| 2021 | AYND1 | K-Nearest Neighbors | 0.21 | 0.04 | 300.89 |
| 2021 | AYND1 | Random Forest | 0.31 | 0.09 | 311.82 |
| 2021 | AYND2 | gBLUP | 0.34 | 0.12 | 235.57 |
| 2021 | AYND2 | Multiple Linear Regression | 0.49 | 0.24 | 219.79 |
| 2021 | AYND2 | K-Nearest Neighbors | 0.39 | 0.15 | 162.04 |
| 2021 | AYND2 | Random Forest | 0.45 | 0.20 | 180.89 |
| 2021 | ELITE | gBLUP | 0.36 | 0.13 | 236.51 |
| 2021 | ELITE | Multiple Linear Regression | 0.56 | 0.31 | 249.92 |
| 2021 | ELITE | K-Nearest Neighbors | 0.32 | 0.10 | 239.97 |
| 2021 | ELITE | Random Forest | 0.55 | 0.30 | 392.09 |

## ABBREVIATIONS

AGL: Above Ground Level
AR1: Autoregressive Order 1
ARDEC: Agricultural Research, Development and Education Center
AYN: Advanced Yield Nursery (F_3:6_)
AYND1: Advanced Doubled Haploid Yield Nursery 1
AYND2: Advanced Doubled Haploid Yield Nursery 2
CSU: Colorado State University
DH: Doubled Haploid
DNA: Deoxyribonucleic Acid
ELITE: Colorado State University Elite Trial (F_3:7+_)
ExGI: Excess Green Index
gBLUP: Genomic Best Linear Unbiased Prediction
GCPs: Ground Control Points
GEBV: Genomic Estimated Breeding Value
GPS: Global Positioning System
GS: Genomic Selection
IID: Identically and Independently Distributed
KNN: K-nearest neighbors
LWIR: Long-Wave Infrared
MLR: Multiple Linear Regression
NDRE: Normalized Difference Red Edge Index
NDVI: Normalized Difference Vegetation Index
NIR: Near Infrared
PPE: Phenomic Estimated Breeding Value
PS: Phenomic Selection
PYN: Preliminary Yield Nursery (F_3:5_)
RF: Random Forest
RGB: Red-Green-Blue Color Spectra
RSME: Residual Square Mean Error
RTK: Real Time Kinematics
S1: Scenario 1
S2: Scenario 2
S3: Scenario 3
TBV: True Breeding Value
UAV: Unoccupied Arial Vehicle

## REFERENCES

Alexandratos, N., & Bruinsma, J. (2012). World agriculture towards 2030/2050: The 2012 revision.

Appels, R., Eversole, K., Feuillet, C., Keller, B., Rogers, J., Stein, N., Pozniak, C. J., Choulet, F., Distelfeld, A., & Poland, J. (2018). Shifting the limits in wheat research and breeding using a fully annotated reference genome. Science, 361(6403), eaar7191.

Berners-Lee, M., Kennelly, C., Watson, R., & Hewitt, C. (2018). Current global food production is sufficient to meet human nutritional needs in 2050 provided there is radical societal adaptation. Elem Sci Anth, 6(1).

Biau, G., & Scornet, E. (2016). A random forest guided tour. TEST, 25(2), 197–227. https://doi.org/10.1007/s11749-016-0481-7

Breiman, L. (2001). Random forests. Machine Learning, 45(1), 5–32.

Browning, B. L., Zhou, Y., & Browning, S. R. (2018). A one-penny imputed genome from next-generation reference panels. The American Journal of Human Genetics, 103(3), 338–348. https://doi.org/10.1016/j.ajhg.2018.07.015

Butler, D., Cullis, B. R., Gilmour, A., & Gogel, B. (2009). ASReml-R reference manual. The State of Queensland, Department of Primary Industries and Fisheries, Brisbane.

Danecek, P., Auton, A., Abecasis, G., Albers, C. A., Banks, E., DePristo, M. A., Handsaker, R. E., Lunter, G., Marth, G. T., Sherry, S. T., McVean, G., Durbin, R., & Genomes Project Analysis, G. (2011). The variant call format and VCFtools. Bioinformatics, 27(15), 2156– 2158. https://doi.org/10.1093/bioinformatics/btr330

Freeman, K. W., Raun, W. R., Johnson, G. V., Mullen, R. W., Stone, M. L., & Solie, J. B. (2003). Late-season Prediction Of Wheat Grain Yield And Grain Protein. Communications in Soil Science and Plant Analysis, 34(13–14), 1837–1852. https://doi.org/10.1081/CSS-120023219

Glaubitz, J. C., Casstevens, T. M., Lu, F., Harriman, J., Elshire, R. J., Sun, Q., & Buckler, E. S. (2014). TASSEL-GBS: a high capacity genotyping by sequencing analysis pipeline. PloS One, 9(2), e90346.

Gutierrez-Gonzalez, J. J., Mascher, M., Poland, J., & Muehlbauer, G. J. (2019). Dense genotyping-by-sequencing linkage maps of two Synthetic W7984×Opata reference populations provide insights into wheat structural diversity. Scientific Reports, 9(1), 1793. https://doi.org/10.1038/s41598-018-38111-3

Hassan, M. A., Yang, M., Rasheed, A., Yang, G., Reynolds, M., Xia, X., Xiao, Y., & He, Z. (2019). A rapid monitoring of NDVI across the wheat growth cycle for grain yield prediction using a multi-spectral UAV platform. Plant Science, 282, 95–103. https://doi.org/10.1016/j.plantsci.2018.10.022

Krause, M. R., González-Pérez, L., Crossa, J., Pérez-Rodríguez, P., Montesinos-López, O., Singh, R. P., Dreisigacker, S., Poland, J., Rutkoski, J., Sorrells, M., & others. (2019). Hyperspectral reflectance-derived relationship matrices for genomic prediction of grain yield in wheat. G3: Genes, Genomes, Genetics, 9(4), 1231–1247.

Kuhn, M. (2008). Building predictive models in R using the caret package. Journal of Statistical Software, 28(1), 1–26.

Kuhn, M. (2022). caret: Classification and Regression Training. https://CRAN.R-project.org/package=caret

Lane, H. M., Murray, S. C., Montesinos-López, O. A., Montesinos-López, A., Crossa, J., Rooney, D. K., Barrero-Farfan, I. D., De La Fuente, G. N., & Morgan, C. L. S. (2020). Phenomic selection and prediction of maize grain yield from near-infrared reflectance spectroscopy of kernels. The Plant Phenome Journal, 3(1), e20002. https://doi.org/10.1002/ppj2.20002

Li, H., & Durbin, R. (2009). Fast and accurate short read alignment with Burrows-Wheeler transform. Bioinformatics, 25(14), 1754–1760. https://doi.org/10.1093/bioinformatics/btp324

Meuwissen, T. H., Hayes, B. J., & Goddard, M. E. (2001). Prediction of total genetic value using genome-wide dense marker maps. Genetics, 157(4), 1819–1829.

Montesinos-López, O. A., Montesinos-López, A., Crossa, J., de Los Campos, G., Alvarado, G., Suchismita, M., Rutkoski, J., González-Pérez, L., & Burgueño, J. (2017). Predicting grain yield using canopy hyperspectral reflectance in wheat breeding data. Plant Methods, 13, 1–23.

R Core Team. (2022). R: A Language and Environment for Statistical Computing. R Foundation for Statistical Computing. https://www.R-project.org/

Ray, D. K., Mueller, N. D., West, P. C., & Foley, J. A. (2013). Yield trends are insufficient to double global crop production by 2050. PloS One, 8(6), e66428.

Rincent, R., Charpentier, J.-P., Faivre-Rampant, P., Paux, E., Le Gouis, J., Bastien, C., & Segura, V. (2018). Phenomic Selection Is a Low-Cost and High-Throughput Method Based on Indirect Predictions: Proof of Concept on Wheat and Poplar. G3 Genes|Genomes|Genetics, 8(12), 3961–3972. https://doi.org/10.1534/g3.118.200760

Robert, P., Brault, C., Rincent, R., & Segura, V. (2022). Phenomic Selection: A New and Efficient Alternative to Genomic Selection Genomic selection (GS). In Genomic Prediction of Complex Traits: Methods and Protocols (pp. 397–420). Springer.

Sandhu, K. S., Merrick, L. F., Sankaran, S., Zhang, Z., & Carter, A. H. (2022). Prospectus of Genomic Selection and Phenomics in Cereal, Legume and Oilseed Breeding Programs. Frontiers in Genetics, 12, 829131. https://doi.org/10.3389/fgene.2021.829131

Santra, M., Wang, H., Seifert, S., & Haley, S. (2017). Doubled haploid laboratory protocol for wheat using wheat–maize wide hybridization. Wheat Biotechnology: Methods and Protocols, 235–249.

